# An International Inter-Laboratory Digital PCR Study Demonstrates High Reproducibility for the Measurement of a Rare Sequence Variant

**DOI:** 10.1101/077917

**Authors:** Alexandra S. Whale, Alison S. Devonshire, George Karlin-Neumann, Jack Regan, Leanne Javier, Simon Cowen, Ana Fernandez-Gonzalez, Gerwyn M. Jones, Nicholas Redshaw, Julia Beck, Andreas W. Berger, Valérie Combaret, Nina Dahl Kjersgaard, Lisa Davis, Frederic Fina, Tim Forshew, Rikke Fredslund Andersen, Silvia Galbiati, Álvaro González Hernández, Charles A. Haynes, Filip Janku, Roger Lacave, Justin Lee, Vilas Mistry, Alexandra Pender, Anne Pradines, Charlotte Proudhon, Lao Saal, Elliot Stieglitz, Bryan Ulrich, Carole A. Foy, Helen Parkes, Svilen Tzonev, Jim F. Huggett

**Author notes:** Corresponding author: Jim Huggett.

## Abstract

This study tested the claim that digital PCR (dPCR) can offer highly reproducible quantitative measurements in disparate labs. Twenty-one laboratories measured four blinded samples containing different quantities of a *KRAS* fragment encoding G12D, an important genetic marker for guiding therapy of certain cancers. This marker is challenging to quantify reproducibly using qPCR or NGS due to the presence of competing wild type sequences and the need for calibration. Using dPCR, eighteen laboratories were able to quantify the G12D marker within 12% of each other in all samples. Three laboratories appeared to measure consistently outlying results; however, proper application of a follow-up analysis recommendation rectified their data. Our findings show that dPCR has demonstrable reproducibility across a large number of laboratories without calibration and could enable the reproducible application of molecular stratification to guide therapy, and potentially for molecular diagnostics.

**SIGNIFICANCE STATEMENT:** The poor reproducibility of molecular diagnostic methods limits their application in part due to the challenges associated with calibration of what are relative measurement approaches. In this study we investigate the performance of one of the only absolute measurement methods available today, digital PCR (dPCR), and demonstrated that when compared across twenty-one laboratories, dPCR has unprecedented reproducibility. These results were achieved when measuring a challenging single nucleotide variant and without calibration to any reference samples. This opens the possibility for dPCR to offer a method to transform reproducibility in the molecular diagnostic field, both by direct use as well as in support of other currently used clinical methods.

## INTRODUCTION

Quantification using real-time quantitative PCR (qPCR) was first described over two decades ago (1, 2). Since then, despite its application to a wide range of preclinical research areas and its capacity for performing precise nucleic acid quantification, there are only a small number of examples where it has been successfully translated into the clinic for quantification, primarily in the fields of clinical virology (3) and management of patients with CML (4). Obstacles to its widespread clinical adoption have been the challenges of standardisation and defining the technical reproducibility of the method (5, 6). In qPCR, reproducibility is often poor due to the fact that while it can be very precise, it can be biased (6); it is not unusual for viral titres to vary by several orders of magnitude between different laboratories in the absence of a calibrator material (7).

Digital PCR (dPCR), where the presence or absence of a target molecule is detected in a binary and absolute fashion, is an alternative method that has the potential to be considerably more reproducible than qPCR. It also offers a number of advantages that include high precision (8–13) and improved sensitivity and specificity (14–18), all without reliance on a calibration curve (19, 20). This high precision has been employed to value- assign certified reference materials that include a plasmid for *BCR-ABL1* monitoring in chronic myeloid leukaemia (21) and genomic DNA for *HER2* amplification detection in breast cancer (22). Prototype reference materials of plasmid, genomic DNA and whole bacteria in synthetic sputum for Tuberculosis have also been evaluated using dPCR (12, 23). However, the method has not yet been demonstrated to surmount the challenges of end-user inter-laboratory reproducibility that will be necessary if it is to be used for routine clinical use.

The central aim of this study was to evaluate the reproducibility of droplet-based dPCR using an international end-user inter-laboratory comparison. The chosen target was a *KRAS* single nucleotide variant (SNV), one of the most challenging types of cancer biomarkers, which is being pursued vigorously in liquid biopsy translational studies that predict non-response to specific therapies in colorectal carcinomas (24, 25) and non-small cell lung cancers (26).

## RESULTS

### Design of the study

Twenty-one participant laboratories were enrolled in the study to measure blinded samples that contained varying fractional abundances of a 186 bp plasmid fragment containing the *KRAS* G12D variant and/or the wild-type (wt) sequence **(Supplementary Document 1** and **Fig. S1).** Each participant laboratory was provided with three units of each of the four blinded test samples. These test samples were at different nominal G12D fractional abundances: 12%, 0.9%, 0.17% and 0% and randomly assigned a sample letter (A-D) **(Table 1).** Additionally two control samples were provided as three units of the negative control (0% G12D; which was the same as sample B and denoted NEG) and a single unit of the positive control (12% G12D; which was the same as sample D and denoted POS). The %G12D of the POS control was not revealed to the participants and so could not be used as a calibrator for the blinded samples. Reagents and QX100/200™ consumables from common manufacturing lots **(Supplementary Document 2)** were provided to the participants along with a full protocol **(Supplementary Document 3),** that included the reaction preparation procedure and the plate layout, to minimise these as possible sources of variation. All packages were distributed by LGC who coordinated the study.

**Table 1.** Values for the *KRAS* target materials obtained during characterisation and inter-laboratory comparison.

| Material |  | Characterisation |  |  | Inter-laboratory participant results<sup>#</sup> |  |  |  |  |
| --- | --- | --- | --- | --- | --- | --- | --- | --- | --- |
| Name | Target | Nominal* | Mean* | U<sub>rel</sub> | Median* | % difference<sup>\$</sup> | MAD<sub>E</sub> | Lower 99% CI | Upper 99% CI |
| A | [WT] | 2000 | 2400 | 1.0% | 2267.00 | 6% | 182.85 | 1795.24 | 2738.76 |
|  | [G12D] | 3 | 4.9 | 5.9% | 4.74 | 3% | 0.59 | 3.23 | 6.26 |
|  | % G12D | 0.17 | 0.20 | 6.0% | 0.21 | 5% | 0.01 | 0.18 | 0.24 |
| B | [WT] | 2000 | 2300 | 1.7% | 2197.00 | 4% | 162.31 | 1781.50 | 2612.50 |
|  | [G12D] | 0 | 0.58 | 16% | 0.42 | 28% | 0.19 | -0.08 | 0.92 |
|  | % G12D | 0.00 | 0.025 | 16% | 0.02 | 20% | 0.01 | 0.00 | 0.04 |
| C | [WT] | 2000 | 2300 | 1.4% | 2273.78 | 1% | 257.71 | 1608.88 | 2938.68 |
|  | [G12D] | 18 | 22 | 4.5% | 22.67 | 3% | 1.78 | 18.08 | 27.26 |
|  | % G12D | 0.90 | 0.96 | 3.5% | 1.01 | 6% | 0.05 | 0.89 | 1.14 |
| D | [WT] | 1760 | 2100 | 1.5% | 1975.00 | 6% | 161.98 | 1557.10 | 2392.90 |
|  | [G12D] | 240 | 310 | 1.6% | 300.80 | 3% | 21.55 | 245.19 | 356.41 |
|  | % G12D | 12.00 | 13.00 | 0.92% | 13.16 | 1% | 0.15 | 12.79 | 13.54 |
| NEG | [WT] | 2000 | 2300 | 1.7% | 2264.67 | 2% | 217.51 | 1707.85 | 2821.48 |
|  | [G12D] | 0 | 0.58 | 16% | 0.49 | 16% | 0.14 | 0.12 | 0.85 |
|  | % G12D | 0.00 | 0.025 | 16% | 0.02 | 19% | 0.01 | -0.01 | 0.05 |
| POS | [WT] | 1760 | 2100 | 1.5% | 1990.00 | 5% | 139.90 | 1629.07 | 2350.93 |
|  | [G12D] | 240 | 310 | 1.6% | 294.67 | 5% | 23.23 | 234.72 | 354.61 |
|  | % G12D | 12.00 | 13.00 | 0.92% | 13.10 | 1% | 0.32 | 12.27 | 13.93 |

In order to remain ambiguous to the participants, the sample name (A-D) was randomly assigned to each of the four materials so that there was no correlation between sample name and G12D fractional abundance. The two control materials, NEG and POS, were generated from the sample materials as Sample D and Sample B, respectively and so have identical characterisation values. \*[G12D] and [WT] are given as copies/ μL in the reaction based on following the study protocol where 8.8 μL of each study material is added to 22 μL reaction during set up; this is the metric participants were asked to report. The nominal concentrations are the approximate values the materials were prepared at. For characterisation of the materials, the mean concentration and the relative expanded uncertainty (U<sub>rel</sub>), that is the equivalent of the 95% confidence interval reported as a percentage of the mean, were calculated from the homogeneity study with triplicate measurements (within-unit) from six units (between-units). All values are given to 2 significant figures. <sup>#</sup>For the inter-laboratory participant results, the values are calculated from the twenty-one original submitted values. <sup>\$</sup>The percentage difference between the mean concentration obtained from the homogeneity study with the participant median value. The MAD<sub>E</sub> is the median absolute deviation.

Participants used their own QX100/200™ Droplet Digital PCR System (Bio-Rad) with manual droplet generation and were requested to analyse their data using their routine procedure. Each laboratory submitted their mean value for the *KRAS* G12D and *wt* copy number concentrations and %G12D with the associated 95% confidence intervals for each sample, control and no template controls (NTCs) in a pre-designed spreadsheet format (summarised in **Table S1**). Details of their analysis method **(Table S2)** and screen shots of the analysis (not shown) were also requested.

### Inter-laboratory comparison

For the quantification of the *KRAS* G12D and *wt* copy number concentrations and %G12D, dPCR measurements from the twenty-one laboratories were highly accurate over the three orders of magnitude (~0.42 to ~2200 copies/µL) reflected in the blinded test samples **(Fig. 1)** and two control samples **(Fig. S2).** There were two groups of results. The majority of the laboratories (18/21) measured both the copy number concentrations and fractional abundance of all samples within 20% of each other. The remaining three laboratories consistently under-quantified the G12D copy number concentration and associated fractional abundance in all samples, though they did report consensus values for the *wt* target concentration. There was no significant difference in the copy number concentrations or fractional abundance of sample B and NEG (p > 0.12) or sample D and POS (p > 0.82). The median participant values for all six samples were within 6% of the mean values obtained in the homogeneity study performed by the co-ordinating laboratory (LGC) that manufactured and characterised the samples (**Table 1**). The exception was for the G12D copy number concentration and fractional abundance of sample B and NEG, that contained no *KRAS* G12D target, that varied by 16-28%.

**Figure 1.**
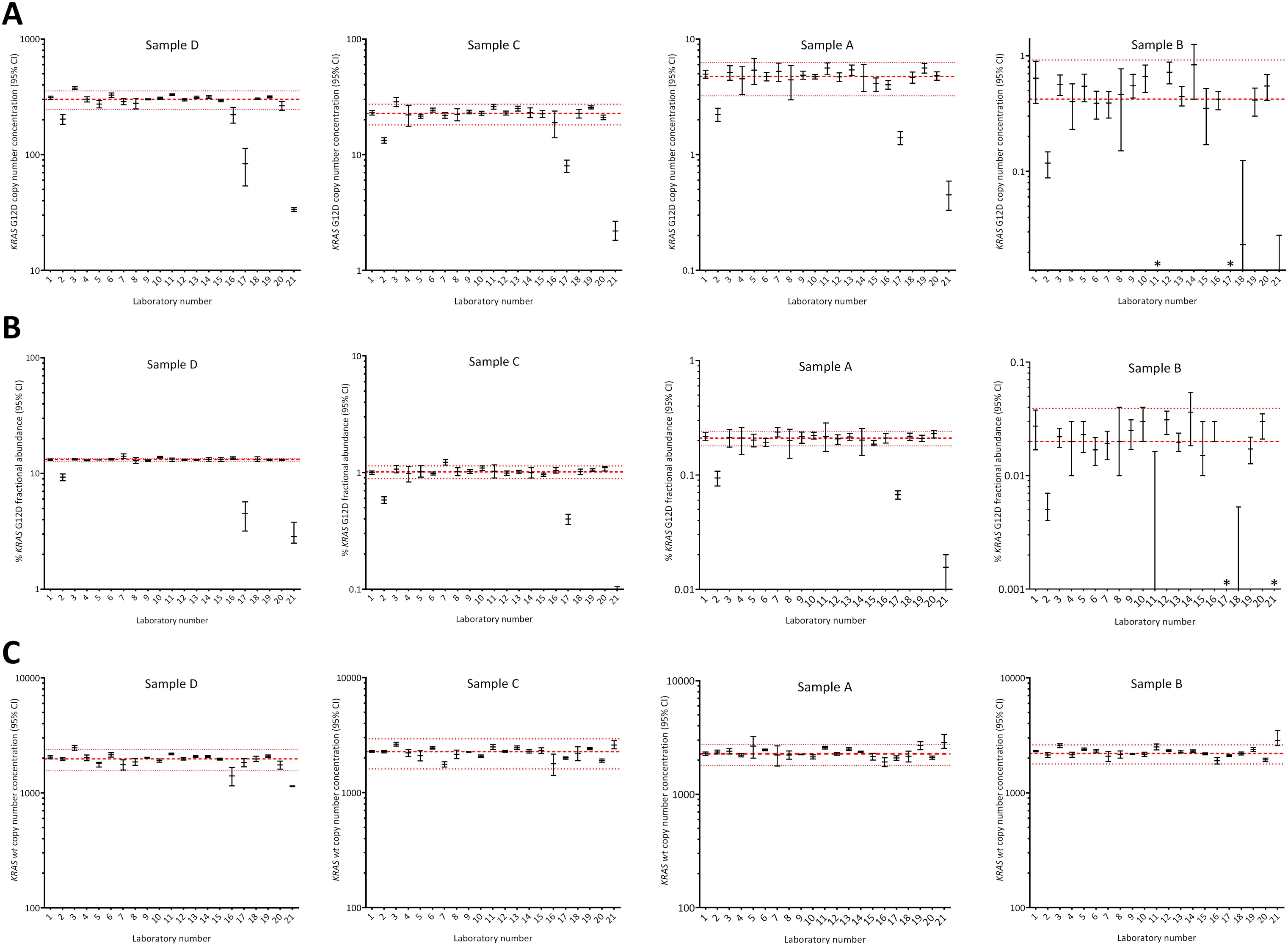
Inter-laboratory comparison. The originally submitted values from all twenty-one laboratories are shown for Samples A-D in descending KRAS G12D copy number concentration from left to right. For each graph the anonymised laboratory number is shown in the x-axis with the (A) KRAS G12D or (C) wt copy number concentration (copies/µL in the reaction) shown on the y-axis as indicated. For the fractional abundance graphs (B), the %G12D is shown on the y-axis. For each participant, the submitted mean value is plotted as a short black horizontal line together with the 95% confidence interval based on triplicate measurements of three units (n=9). The red horizontal dashed line represents the median value across labs and the red horizontal dotted lines represent the upper and lower MAD_E_ intervals with 99% confidence. For Sample B, the lower confidence interval is not shown as it is approximately zero. Asterisks (*) just above the x-axis indicate laboratories that reported either a zero value or values below the range of the y-axis. All graphs show two orders of magnitude, shown on the log_10_ scale, though the range varies according to the sample.

Of the first group of results, fourteen laboratories had no outlier values in any of the test or control samples (**Fig. 1** and **S2).** The remaining four laboratories (3, 7, 10 and 16) had outlier values in one or more of the measurements for one or more samples which were associated with moderately higher or lower *wt* or G12D copy number concentration or fractional abundance (>1.5-fold different from the median value). The group that consistently under-quantified the G12D molecules (2, 17 and 21) were all identified as outliers and submitted values that were between 2- and 9-fold lower than the median for the G12D (but not *wt*) copy number concentration and fractional abundance.

### Factors affecting reproducibility

Following visual inspection of the two-dimensional (2D) scatter plots produced by the QuantaSoft ddPCR analysis software, it was hypothesised that the cause of the G12D under-quantification in laboratories 2, 17 and 21 was misclassification of the droplets **(Fig. S3).** This hypothesis was tested by preparation of guidelines for droplet classification that were circulated to all participants **(Supplementary Document 4).** At this stage all participants remained blind to the copy number concentrations. The seven laboratories with one or more outlier were invited optionally to reanalyse their original data following the guidelines. In parallel, the coordinating laboratory reanalysed the seven laboratory data sets following the guidelines.

Four laboratories resubmitted their values (2, 10, 17 and 21) **(Table S3)**. In contrast to their originally submitted values, for two of the laboratories (2 and 21), no significant difference was observed between their resubmitted values and the median values from the original inter-laboratory comparison (p >0.35) suggesting that the guidelines rectified the droplet misclassification **(Fig. 2** and **S4).** However, laboratory 17 resubmitted values that were ~1.2-fold higher than their original submission, but still significantly lower (>2-fold) than the median values from the original inter-laboratory analysis (p ≤0.012) **(Fig. 2** and **S4).** Reanalysis of this data set by the coordinating laboratory identified no significant difference between their values and the median values from the original inter-laboratory analysis (p >0.14) **(Fig. 2** and **S4)** suggesting that droplet misclassification, and not the actual data generated, was still responsible for the under-quantification of this laboratory. Laboratory 10, which was an outlier in only one value in one test sample (the G12D fractional abundance in sample D), resubmitted values that were not significantly different from those obtained by the coordinating laboratory **(Fig. 2** and **S4)** indicating that droplet misclassification was not responsible for this single outlier result. The other three laboratories (3, 7 and 16) communicated that they were satisfied that their original analysis aligned with the guidelines; there was no significant difference between their original values and those obtained by the coordinating laboratory (p >0.34).

**Figure 2.**
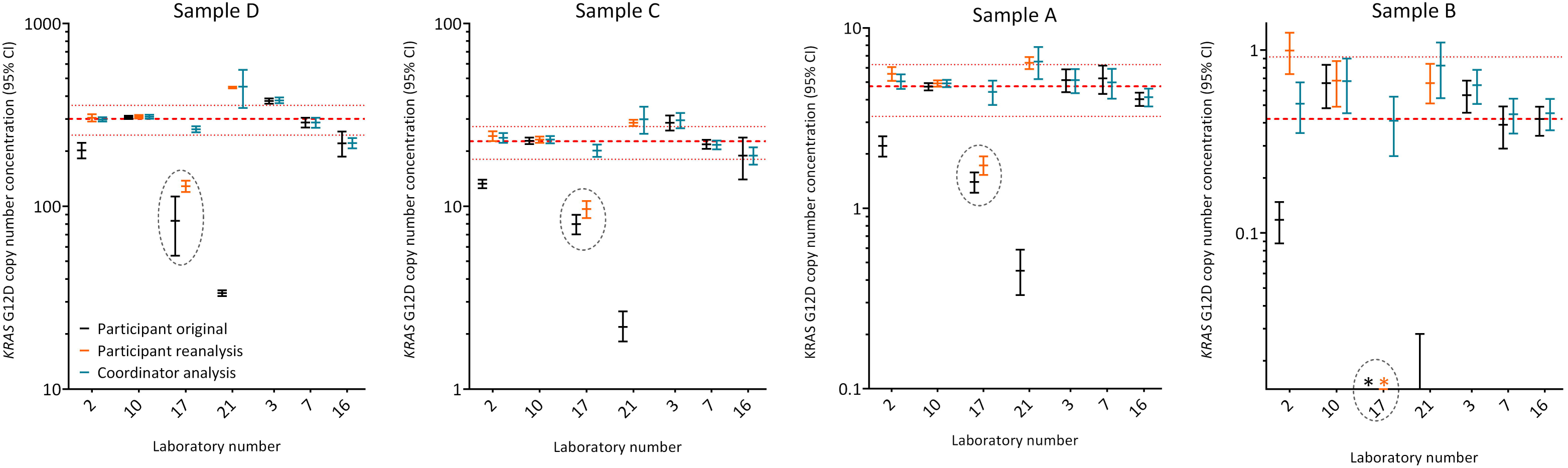
Quantification of *KRAS* G12D following threshold setting guidelines. For each sample the comparison between the copy number concentrations originally submitted (black bar) with the resubmitted values (orange bar) and the coordinator analysis (teal bar) is shown. The red horizontal dashed line represents the median from the *original* data set with the associated red horizontal dotted lines representing the upper and lower MAD_E_ intervals with 99% confidence. The scales are the same as those in Figure 1. Four of the laboratories reanalysed their data (2, 10, 17 and 21) and the remaining three laboratories were satisfied with their original submission (3, 7 and 16) and so do not contain data points for resubmission. The coordinating laboratory analysed the data sets from the seven laboratories. The dashed ovals highlight laboratory 17 that were still under-quantifying the G12D concentration in the four samples compared with the median; while the analysis by the coordinating laboratory did not.

Between-laboratory precision was calculated for all samples and controls using the four resubmitted laboratory values along with the original values from the other seventeen laboratories. For nominal copy number concentrations >0.1 copies/µL, the precision was high across all the samples and targets; CVs were between 7.39% and 11.39% **(Table 2).** Calculation of the theoretical CV (tCV) for each value for each sample showed that the precision based on Poisson alone was generally only slightly smaller than the between-laboratory precision for the G12D and *wt* copy number concentrations in the samples that contained G12D molecules **(Table 2)** indicating a minimum of experimental error. For the corresponding G12D fractional abundances, the precision was high for all samples that contained G12D fragments (CV <4.8%).

**Table 2.** Precision of the inter-laboratory comparison following resubmission.

| Material Name | Target | Recalculated Median* | % difference<sup>§</sup> | CV between labs | tCV |
| --- | --- | --- | --- | --- | --- |
| A | [WT] | 2267.00 | 6% | 8.07% | 1% |
|  | [G12D] | 4.87 | 1% | 11.39% | 10% |
|  | % G12D | 0.21 | 5% | 4.22% |  |
| B | [WT] | 2197.00 | 4% | 7.39% | 1% |
|  | [G12D] | 0.46 | 21% | 34.50% | 35% |
|  | % G12D | 0.02 | 20% | 37.08% |  |
| C | [WT] | 2274.00 | 1% | 11.35% | 1% |
|  | [G12D] | 22.98 | 4% | 7.87% | 5% |
|  | % G12D | 1.02 | 6% | 4.74% |  |
| D | [WT] | 1982.00 | 6% | 7.65% | 1% |
|  | [G12D] | 303.30 | 2% | 5.88% | 1% |
|  | % G12D | 13.20 | 2% | 1.12% |  |
| NEG | [WT] | 2264.67 | 2% | 9.60% | 1% |
|  | [G12D] | 0.54 | 7% | 24.72% | 32% |
|  | % G12D | 0.02 | 20% | 31.26% |  |
| POS | [WT] | 1990.00 | 5% | 6.33% | 1% |
|  | [G12D] | 298.00 | 4% | 7.30% | 1% |
|  | % G12D | 13.24 | 2% | 2.69% |  |

\*[G12D] and [WT] are given as copies/ μL in the reaction as described in Table 1. The values presented in this table are calculated from the four resubmitted values and remaining seventeen original values. <sup>§</sup>The percentage difference between the mean concentration obtained from the homogeneity study (Table 1) with the recalculated median. The tCV is calculated based on the equation described in Devonshire et al. 2015.

For the *wt* only samples, that contained ~40,000 *wt* copies/reaction and no G12D fragments (Samples B and NEG), the identification of positive partitions for the G12D assay represented the false positive rate (FPR) of the assay. A median FPR of 0.02% G12D was observed in these samples that corresponded to a median of ~6 G12D-positive partitions/reaction (minimum of 0 and maximum of 14 across all 21 participants NEG reactions; n=198) **(Table S4)**. Measurements of the *KRAS* G12D molecule in these samples had reduced precision (CV >24%) as would be expected for such low measured values. For the no template controls (NTCs), a median and mode of 0 G12D-positive partitions/reaction was observed (maximum of 3 across all 21 participants, n=186) **(Table S5)** indicating low, if any, contamination in the dPCR reactions.

The *KRAS wt* copy number concentrations were approximately equal for samples A, B and C **(Table 1).** To generate a metric to measure the impact of various deviations from the study protocol, the ratio of the *KRAS wt* copy number concentration of samples C and A was used. From the results of the questionnaire relating to the study protocol **(Table S2)** no significant difference (p >0.14) was observed related to the QX system or version of the QuantaSoft analysis software used, number of times the samples were thawed prior to analysis or any other deviations from the protocol such as the supplier of pipettes, plate sealer or PCR cycler **(Fig. S5).**

Characterisation of each study material established that the between-unit variation did not exceed the within-unit variation (p >0.24 for all study materials) **(Fig. S6)** and so estimated standard uncertainties due to possible inhomogeneity were calculated using the between-unit variation. The relative standard uncertainties were <6% for samples containing G12D molecules **(Table 1)** and so it was concluded that the unit homogeneity of all the study materials was acceptable.

Stability studies identified no significant time or temperature effect over the short term for the G12D and *wt* copy number concentrations with the exception of sample D/POS that had a 1.1-fold decrease in both G12D and *wt* copy number concentrations (but no effect on the fractional abundance, data not shown) when stored on dry ice for 7 days **(Fig. S7).** For the long term stability study, small differences in the G12D and *wt* copy number concentrations were observed over the six month duration of the stability study **(Fig. S8).** For both stability studies the differences were minor (<12%), non-directional (with the exception of sample D/POS), and all participants performed their experiments within one month of receiving the samples, it was concluded that the stability of the materials was satisfactory.

## DISCUSSION

The aim of this study was to investigate whether dPCR could offer reproducible quantification between laboratories in the absence of calibration using a droplet-based technology. We used an international inter-laboratory study focusing on the quantification of a range of SNV concentrations, of varying fractional abundances, using droplet dPCR. Participant laboratories were recruited from a range of institutes including hospitals, universities and research industries. To our knowledge, this is the first study of its kind and could form the basis of a framework for other studies investigating the reproducibility and robustness of a given dPCR method. For this purpose, we used a clinically relevant prognostic model of a *KRAS* point mutation that needs to be determined prior to selection of treatment of specific cancers (27, 28).

We chose a range of G12D to *wt* fractional abundances, several of which are challenging to quantify with currently used analytical methods (29, 30). Furthermore, as the interest in liquid biopsies continues to rise, we directed the selection of the concentration and size of the molecules quantified in this study to match this template type (31, 32). Comparison of the independent measurements from twenty-one laboratories demonstrated the high level of reproducibility of dPCR, defined as a CV of <12%. Further investigation identified that most of this variation could be attributed to deviations in tube-to-tube unit homogeneity and stability of the test materials as well as the Poisson error, thereby rendering this reproducibility all the more impressive.

This study clearly demonstrated that fractional abundancies down to ~0.2% can be reproducibly quantified using dPCR. As the observed FPR was ~0.02%, it suggests that a lower limit of detection of the G12D molecule, such as 0.1%, may be achievable with dPCR. However, the observed ~0.2% fractional abundance was enabled, in part, by the addition of ~40,000 *KRAS* copies per reaction; this level of sensitivity may not always be achievable using *in vivo* extracts such as cfDNA from plasma, that typically yields ~500 copies/mL plasma (33). In addition to cancer models, other rare SNV models would also be applicable to this framework (14, 15) as well as simpler targets that do not require specific measurement of a given variant in the presence of the *wt* allele, such as quantification of specific pathogens or gene fusions (34, 35).

Two distinct aspects of this technique demonstrate that dPCR is reproducible for absolute quantification measurements. Firstly, all participant laboratories were blind to the target quantities of each test sample, yet twenty laboratories were able to correctly quantify all four test samples. Secondly, all the measurements were performed without any form of calibration to a material of known quantity; the materials were simply quantified and results reported directly.

In order to determine sources of discrepancy where they arose, we investigated a number of parameters that could influence reproducibility and robustness. Deviations from the study protocol, such as droplet handling using different pipettes and tips to those recommended did not have significant effects on the quantification. Furthermore, participants used different software versions to collect their data. Versions 1.3 and below use a 0.91 nL droplet volume that has been identified as a source of bias in other studies (36), whilst the smaller volume of 0.85 nL is used in versions 1.6 and above. While the use of different partition volumes can impact on accuracy, in this study, the two partition volumes used did not appear to impact on the between-laboratory reproducibility **(Fig.** S5), suggesting that technical error masked this potential source of bias.

All participants used the same assays, reagents and consumables in this study; another study where these were varied demonstrated good reproducibility (12). Additionally, this study did not evaluate pre-analytical steps such as the extraction, however, with careful optimisation, quantification can be reproducible even when the extraction step is incorporated (23).

Incorrect classification of the positive and negative droplets was a main cause of under-quantification by some laboratories. Partition misclassification has been highlighted as a source of bias in dPCR quantification (37) and can be complicated by a number of factors that include template type, source and matrix in addition to assay and reagent choice and thermocycling conditions (12). When measuring SNV quantities further factors that can impact on correct partition classification include assay specificity, partition specific competition (PSC) and assay FPR. Assay specificity and PSC can make it more difficult to classify the four cluster types (38) while the FPR, if not determined correctly can impact on the sensitivity of the assay and lead to an underestimation.

The presence of G12D false-positive droplets in the wt-only (NEG) control in two of the participant laboratories directed the analysts to be stringent with their droplet classification and opt for increased specificity over sensitivity; such a method has been adopted in another study (39). While this reduced the FPR (increasing specificity), it was at the expense of sensitivity and resulted in a 2 to 3-fold negative bias due to the high total DNA input per reaction. Therefore, for the most sensitive measurements, characterisation of the FPR is needed. However, in order for the clinical utility of this method to be realised, rigorous criteria for calling positive results, along with detailed investigation of the limits of detection and quantification need to be developed and evaluated as illustrated with *EGFR* mutations in lung cancer (40, 41).

## CONCLUSIONS

This study has confirmed that dPCR can perform highly reproducible absolute quantification of an SNV between laboratories differing in their reported value by <12%. This was achieved without calibration, which is not possible using conventional qPCR when measuring samples that contain challenging fractional abundances of <1%. Clinical quantification of SNVs has been proposed for a range of applications such as monitoring donor organ rejection (17) or in the treatment of cancer patients (16) and will likely be needed for the development of precision/stratified medicine to be fully realised. These findings suggest dPCR will have an important role in enabling such measurements. In the short term, this study will contribute to a growing body of evidence that demonstrate that dPCR can offer a valuable method to ensure reproducible quantification of nucleic acids.

## MATERIALS AND METHODS

The coordinating laboratory for this study was LGC (UK). All experiments pertaining to the production and characterisation of the study materials were performed at LGC. All kits and instruments were used following the manufacturer’s instructions.

### Production of the study materials

A portion of the human *KRAS* gene encoding exon 2 and its flanking introns was downloaded from the NCBI database (NCBI accession: NG_007524.1, bases 9788 to 11411). The location of the drug resistant mutation (c. 35G>A) that encodes the amino acid substitution G12D was identified. The inserts were synthesised using overlapping synthetic oligonucleotides and then blunt-end cloned into the EcoRV restriction site of the pUC57 plasmid (Eurofins) to generate two constructs: one containing the wild type (*wt*) sequence and one containing the G12D mutation. Sequence validation was performed by Sanger sequencing on both strands (Eurofins) (**Fig. S9A** and **S9B**).

Ten double restriction digest reactions were set up for each *KRAS* construct each containing approximately 10 µg of DNA with 25 units each of *Afl*III and *NsiI* (New England Biolabs) in a 50 µL reaction volume. The reactions were incubated at 37 °C for three hours and subjected to electrophoresis through a 3% agarose gel. Each 186 bp fragment was identified using the 1kb+ DNA ladder (Invitrogen), gel purified using the Gel Extraction kit (Qiagen) and eluted in 30 µL of EB buffer; the 10 replicate reactions were pooled to give a total eluent volume of 300 µL. The two 186 bp fragments (KRAS *wt* and G12D)were assessed for size and purity using the 2100 Bioanalyzer with the DNA 7500 series II assay (Agilent) (**Fig. S9C** and **S9D**) and the concentration was estimated using the Qubit^®^ 2.0 fluorometer with the dsDNA HS Assay Kit (Invitrogen) to give the ‘nominal’ stated values for each test sample and control. The copy number concentration was estimated based on the molecular weight of the 186 bp fragment (approximately 116 kDa) and the Avogadro constant using a standard method (42); 1 ng corresponds to approximately 2.01 x 10^8^ copies.

The four study materials were manufactured in 50 mL tubes by gravimetrically diluting the *KRAS wt* and G12D purified 186 bp fragments in non-human carrier to their nominal concentrations **(Table 1).** Carrier was commercially available sonicated salmon sperm genomic DNA (Ambion) at a final concentration of 25 ng/µL in the study materials. The study materials were mixed by rotation at 50 rpm at 4 °C for 30 minutes. One test material was prepared from each study material; these were randomly labelled as samples A, B, C or D to reduce analyst bias. In addition, two control materials were prepared from two of the study materials; the negative control (NEG) was the same as Sample B and the positive control (POS) was the same as Sample D. Following mixing, the four test and two control samples were aliquoted (>100 µL) in pre-labelled 0.5 mL tubes (P/N E1405-2120; StarLabs) to generate between 75 and 300 units and stored at -20 °C. To reduce contamination risks, the test materials were manufactured in order of increasing G12D concentration.

### Characterisation of study materials

Unit homogeneity was evaluated across twelve randomly selected units for each study material that were maintained at -20 °C. The short term stability (STS) study was set up isochronously at a range of temperatures to monitor the stability of the study materials during shipment to the participants and the accelerated stability at high temperature. Three units of each of the four study materials were incubated on dry ice (shipment temperature), at 4 °C and 28 °C (accelerated stability temperature) for 2 and 7 days and compared to the baseline temperature (three units maintained at -20 °C). Following all temperature and time point conditions, the study materials were incubated at -20 °C overnight before dPCR analysis. The long term stability (LTS) study was performed to establish the stability of the study materials over the full course of the study. Three units of each of the four study materials were maintained on dry ice for 5 days to mimic shipping conditions, before being transferred to -20 °C before storage for 1 or 7 months prior to dPCR analysis; parallel experiments were compared with samples maintained at -20 °C for the duration of the LTS. Details of the dPCR analysis performed are in the relevant section below.

### Droplet Digital PCR

All dPCR experiments were implemented in accordance with the dMIQE guidelines **(Table S6)** (20) using the QX100™ or QX200™ Droplet Digital PCR System (Bio-Rad). Briefly, 22 µL reactions were established containing ddPCR Supermix for Probes with no dUTP, PrimePCR™ ddPCR™ Mutation Assays for KRAS p.G12D and KRAS WT (assay details in the relevant section below), 8.8 µL of study material and made up to volume with nuclease-free water. No template controls (NTCs) were performed using nuclease-free water in place of template in every experiment. Droplets generation was performed manually (not with the AutoDG™ system) from 20 µL of the reaction following manufactures guidelines. Thermocycling conditions were 95 °C for 10 minutes, 40 cycles of 94 °C for 30 seconds and 55 °C for 1 minute, followed by 98 °C for 10 minutes and a 4 °C hold. The ramp rate for each step was set to 2 °C/second. Droplet reading was performed following manufactures guidelines. Details of the protocol are given in **Supplementary Document 3.**

For the study material characterisation, experiments were performed using the QX200™ droplet digital PCR system with a randomised plate layout. Every study unit was analysed with triplicate wells. Aliquots of the carrier the study materials were made up with were also evaluated for the presence of *KRAS* sequences (carrier only controls); no positive droplets for the *KRAS* G12D were observed. If multiple 96-well plates were necessary, replicate units were distributed between the different plates with the triplicate dPCR wells for each unit located on the same plate. The commercially available PrimePCR™ ddPCR™ Mutation Assays used were KRAS p.G12D, Human (P/N 10031246, Assay ID dHsaCP2000001) and KRAS WT for p.G12D, Human (P/N 10031249, Assay ID dHsaCP2000002) that generate a 90 nt amplicon. All PrimePCR assays are designed and validated by the manufacturer on the QX100/200 droplet digital PCR system. The QuantaSoft software version 1.7.4.0917 was used to collect the data in the ‘ABS’ mode. The status of each well was checked and wells containing <10,000 droplets or a baseline drop in the double-negative droplets compared with the NTC wells were omitted from the data set. Thresholds were set using the ‘auto analyze’ and ‘combined wells’ setting in the 2D amplitude mode to define the positive and negative droplets. The data was exported as a .csv file for further analysis.

Prior to the inter-laboratory comparison, a pilot study was performed to test the international shipment method of the study materials, reagents and consumables, and the clarity of the protocol. Three Bio-Rad laboratories (USA, France and Germany) participated in the pilot study to confirm that the shipment conditions were acceptable. Following the data analysis, the preparation protocol for the reagents and the dPCR protocol were amended. The final documents and details are included in the supplementary information as cited below.

Participant laboratories were provided with the protocol by the coordinating laboratory **(Supplementary Document 3).** The commercially available PrimePCR™ ddPCR™ Mutation Assays used were KRAS p.G12D, Human (P/N 10031246, Assay ID **dHsaCP2500596**) and KRAS WT for p.G12D, Human (P/N 10031249, Assay ID dHsaCP2500597) that generates a smaller amplicon of 57 nt for each assay compared with the assay used for material characterisation. Each unit was analysed in triplicate wells alongside eight no template controls (NTCs) to give a total of 56 reaction wells in a single experiment; a plate layout was provided in the protocol. The droplets were read in the ‘ABS’ mode using a QuantaSoft template file provided by the coordinating laboratory. Participants could use either the QX100™ or QX200™ Droplet Digital™ system. Each laboratory was asked to complete a questionnaire with details of their ddPCR instrumentation, software version and data analysis methods (Supplementary Document 3). They were requested to report the concentration (copies/µL in the reaction) and percentage allelic frequency of the *KRAS* G12D in each of the four test samples and two controls with the associated 95% confidence intervals as well as the result of their NTCs. In addition to the questionnaire, laboratories were asked to submit their raw data (Quantasoft .csv and *.qlp* files) and screen shots of the 2D scatter plots of the control samples (NEG, POS and NTC).

### Data analysis

For the characterisation of the test materials, exported .csv files were imported into MS Excel 2010 in the first instance with further analysis performed in the R statistical programming environment version 3.1.1 (http://www.r-project.org/) and Prism 6 (GraphPad). A tab deliminated file was generated in MS Excel 2010 containing the well number, sample name, unit number, replicate well number, software generated copies/µL in the reaction for the *KRAS* G12D and *wt* molecule.

For assessment of unit homogeneity, ANOVA and mixed effects models with maximum likelihood estimation described previously (43) were used to assess plate and unit variation, and to estimate the between-unit standard deviation. Analysis of STS and LTS study was performed by fitting a linear model to each test material with time and temperature as the covariates to determine if the copy number concentration varied with storage time; the time variate was treated as a continuous variable. Statistical significance was identified using the Bonfernoni correction for false discovery rate (44) so that significance was identified when *p* <0.004 for the STS and *p* <0.01 for the LTS.

The reported values and confidence intervals from each end-user laboratory were transcribed into Prism 6 for analysis and generation of graphs. The data was tested for normality using the Shapiro-Wilk test with non-normality assigned to data when *p* <0.05. Robust statistics were used for the non-normally distributed data to estimate the consensus values and the between lab dispersion using the median and median absolute deviation (MAD), respectively. The MAD was calculated as follows using equation [1]:

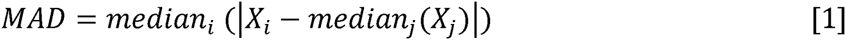

Where *median*_*i*_ is the median of the reported values from the laboratories, and *median*_*j*_ is the median of the absolute difference between the reported values and median. The MAD was then used to estimate the MAD_E_ using equation [2]:

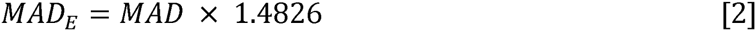

Where the 1.4826 was selected as the consistency constant; for details on the calculation of this value see Staudt *et al.,* 1990 (45). Submitted values outside the upper and lower 99% confidence intervals (calculated using 2.58 MAD_E_ distributions away from the median) were identified as outliers; this level of confidence was chosen as there were more than 20 laboratories enrolled in the study. A robust estimate of reproducibility that down weighs the influence of outliers was calculated using equation [3]:

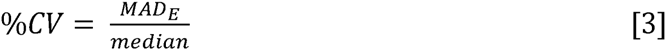

Paired t-tests were used to compare two groups of data; for example, to test for differences between sample B and the negative control, or between the participant resubmitted data and the coordinating laboratory analysis.

## ACKNOWLEDGEMENTS

The work described in this paper was funded by the UK government Department for Business, Energy & Industrial Strategy (BEIS) and the European Metrology Research Programme (EMRP) joint research project (SIB54) Bio-SITrace (http://biositrace.lgcgroup.com) which is jointly funded by the EMRP participating countries within EURAMET and the European Union. The authors would like to thank David Littlemore (LGC) for his central support in the distribution of the materials needed for this study and Pia Scheu (Bio-Rad, Munich, Germany) and Ronald Lebofsky (Bio-Rad, Marnes, France) for their assistance in running pilot experiments to test shipping of samples and kits, and the clarity of the protocol prior to the end-user inter-laboratory study. The authors would also like to thank Cloud Paweletz (Dana Farber Cancer Institute) for critical review of the manuscript prior to submission.

The following authors provided technical and analytical support to the participant laboratories: Isabelle Iacono and Stéphanie Brejon (Centre Léon Berard), Ekkehard Schuetz (Chronix Biomedical), Ana Patiño-Garcia (Clínica Universidad de Navarra), Antoine Carlioz (Assistance Publique Hôpitaux de Marseille), Reinhold Pollner (Genoptix), Virginie Poulot (Tenon Hospital), Anne Casanova and Gilles Favre (Institut Claudius Regaud), Charles Decraene (Institut Curie), Ben Ho Park (Johns Hopkins), Anthony M. George (Lund University), Peter Kaltoft Böhm Nielsen (Sjællands Universitetshospital), Vito Lampasona (Division of Genetics and Cell Biology, and Division of Genetics and Cell Biology & Diabetes Research Institute, San Raffaele Scientific Institute), Julian Downward (Institute of Cancer Research), Daniel Schwerdel (Ulm University), Alice Gutteridge (University College London), Curtis B. Hughesman (University of British Columbia), Daniel Fernandez Garcia and Jacqueline A. Shaw (University of Leicester) and Helen Huang (MD Anderson Cancer Center). The Dana Farber Cancer Institute would also like to acknowledge the Expect Miracles Foundation.

## AUTHOR CONTRIBUTIONS

A.S.W., A.S.D., G.K-N., J.R., S.C., S.T and J.F.H. designed the study, and analysed and interpreted the data. A.S.W., A.F-G., G.M.J. and N.R. produced and characterised the material. L.J. prepared and performed the quality control on the reagents and assays. G.K-N, J.F.H. and S.T. conceived the study. C.A.F., H.P. and J.F.H. obtained funding for the study. J.B., A.W.B., V.C., N.D.K., L.D., F.F., T.F., R.F.A., S.G., A.G.H., C.A.H., F.J., R.L., J.L., V.M., A.P., A.P., C.P., L.S., E.S. and B.U performed the experiments for the inter-lab study. The manuscript was written by A.S.W., G.K-N., and J.F.H. All authors reviewed and provided editorial comments on the manuscript.

